# iCellR: Combined Coverage Correction and Principal Component Alignment for Batch Alignment in Single-Cell Sequencing Analysis

**DOI:** 10.1101/2020.03.31.019109

**Authors:** Alireza Khodadadi-Jamayran, Joseph Pucella, Hua Zhou, Nicole Doudican, John Carucci, Adriana Heguy, Boris Reizis, Aristotelis Tsirigos

## Abstract

Under-sampling RNA molecules and low-coverage sequencing in some single cell sequencing technologies introduce zero counts (also known as drop-outs) into the expression matrices. This issue may complicate the processes of dimensionality reduction and clustering, often forcing distinct cell types to falsely resemble one another, while eliminating subtle, but important differences. Considering the wide range in drop-out rates from different sequencing technologies, it can also affect the analysis at the time of batch/sample alignment and other downstream analyses. Therefore, generating an additional harmonized gene expression matrix is important. To address this, we introduce two separate batch alignment methods: Combined Coverage Correction Alignment (CCCA) and Combined Principal Component Alignment (CPCA). The first method uses a coverage correction approach (analogous to imputation) in a combined or joint fashion between multiple samples for batch alignment, while also correcting for drop-outs in a harmonious way. The second method (CPCA) skips the coverage correction step and uses k nearest neighbors (KNN) for aligning the PCs from the nearest neighboring cells in multiple samples. Our results of nine scRNA-seq PBMC samples from different batches and technologies shows the effectiveness of both these methods. All of our algorithms are implemented in R, deposited into CRAN, and available in the iCellR package.

## INTRODUCTION

Single-cell sequencing of transcripts (RNAs) has become extremely popular in recent years [1], and many large-scale projects such as Human Cell Atlas (HCA) [2] and Mouse Cell Atlas (MCA) [3] are examples of accelerating growth. While many labs produce such data, integrating these data and correcting for batch differences has emerged as a crucial step in the process. Thus, researchers have developed a variety of methods, such as Mutual Nearest Neighbors (MNN) [4], Batch Balanced K Nearest Neighbors (BBKNN) [5], Harmony [6] and Seurat Multiple Canonical Correlation Analysis (MultiCCA) [7], to correct batch effects by aligning different samples.

In addition, single-cell sequencing technologies suffer from low capture rates or under-sampling of RNA molecules, also known as drop-outs [8]. Different sequencing technologies have varying drop-out and gene coverage rates. Although, some labs have developed methods, such as Markov Affinity-based Graph Imputation of Cells (MAGIC) [8], DrImpute [9], Variability-preserving ImPutation for Expression Recovery (VIPER) [10] and scImpute [11], none have developed a method to simultaneously harmonize the drop-out and gene converge rates across different technologies and batches.

Here, we introduce a method to solve both problems in one step. Our Combined Coverage Correction Alignment (CCCA) algorithm not only performs an accurate batch alignment but also harmonizes the gene coverage and drop-out rates. Our Coverage Correction (CC) algorithm is analogous to imputation and performs with a higher accuracy than the other methods mentioned above. Furthermore, we also introduce Combined Principal Component Alignment (CPCA), which skips coverage correction and employs principle components for alignment instead of imputed expression matrix.

## METHODS

### Coverage Correction (CC)

To fill the drop-outs using this method, we perform a Principle Component Analysis (PCA) on the expression matrix and then calculate the distances (Euclidian by default) between the cells using the first few PCs (10 PCs by default). Then, we use KNN [12] to find the nearest neighboring cells (k=10 by default) per each cell (root cell). We then average the expression values of neighboring cells and apply the averaged values to the root cell. This process is repeated for each single cell, creating a high-resolution expression smoothing. Unlike the current imputation methods where the resulted matrices have multiple layers of data transformation (scaling, log, etc.), coverage correction has expression values very close to the ones in the original matrix. This means that fold changes remain similar (in the concept of differential expression analysis) and the data follows a similar pattern. This can be useful if one chooses the option of using coverage corrected data for differential expression and downstream analyses.

### Combined Coverage Correction Alignment (CCCA)

Here, we use coverage correction as an alignment technique and a coverage harmonization method. To do this, we perform a Principle Component Analysis (PCA) on the expression matrix and then calculate the distances (Euclidian by default) between the cells using the first few PCs (30 PCs by default). Then, we collect ‘equal’ numbers of nearest neighboring cells (k=10 by default) for each cell (root cells) ‘from every batch’ and then average the expression values and apply them to the root cell. This is repeated for all the cells. In this process, every cell is considered a root cell only once. This method results in a harmonization among the expression values and gene coverages from different batches and ensures that the drop-out rates are much improved.

### Combined Principle Component Alignment (CPCA)

This method is similar to CCCA; however, it skips the coverage correction part and instead aligns the Principle Components (PCs). In the CPCA method, the expression matrix is replaced with the PC matrix, and PCs are averaged instead of the expression values. CPCA is faster than CCCA, and the batch alignment results are similar. However, CPCA will not create a coverage corrected matrix.

## RESULTS

### Coverage Correction performs better than Markov Affinity-based Graph Imputation of Cells and a few other imputation methods

MAGIC (Markov Affinity-based Graph Imputation of Cells) is a widely used imputation method which employs diffusion data to denoise the expression matrix by filling the drop-outs [8]. To compare CC with MAGIC, we used a commonly analyzed PBMC sample dataset provided by 10x Genomics. Comparing heatmaps of gene expression matrices showed denoising for drop-outs with MAGIC introduced additional noise in a way that the resulting matrix does not show similar expression patterns seen in the original data (Figure 1). Furthermore, MAGIC failed to accurately measure a true gene-gene correlation. For instance, NKG7 and GNLY, two highly correlated genes, are expressed in NK cells and CD8-positive cells and not the other cell types. This pattern can be seen in the original data which includes drop-outs and is much improved using CC, but after MAGIC imputation CD8-positive cells seem to be mixed with other cells (Figure 2). Because the expression matrices imputed with MAGIC undergo a few levels of data transformation, such matrices cannot be used to calculate true fold changes in the concept of differential expression analysis. Our results also show that coverage corrected data, if used for clustering and dimensionality reduction, can illuminate more details by separating more cell types and clusters. For instance, it can separate CD8-positive T cells with distinctive distance from CD4-positive T cells (Figure 2). A few other imputation methods were also compared to CC and while some performed better than others CC was most consistent in following the patterns seen in the original data (Figure S2).

**Figure 1.**
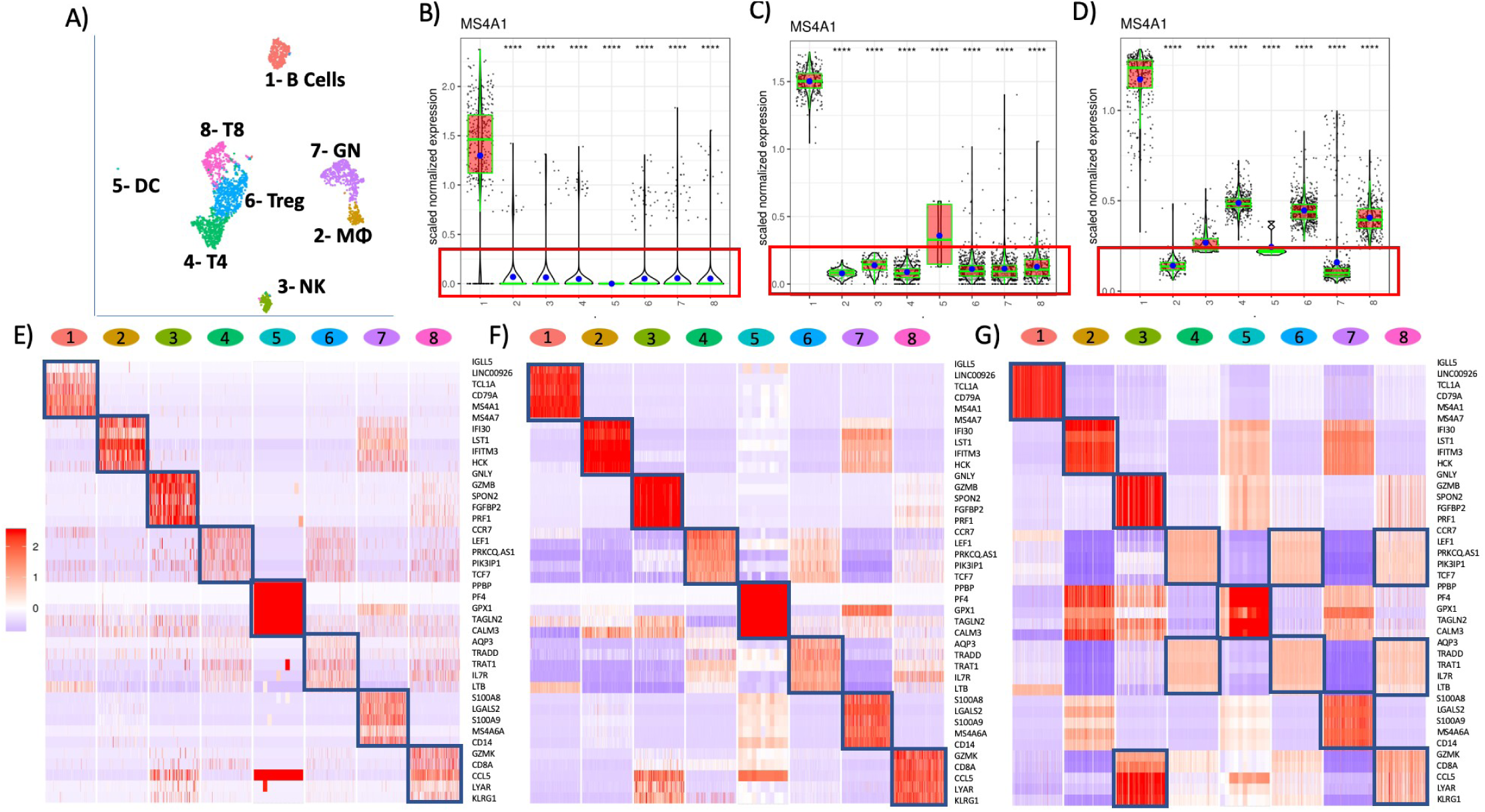
A) UMAP plot of a sample PBMC data showing 8 clusters/cell types. B) Boxplot showing the expression of MS4A1 in the raw expression matrix. Big blue dots in the box plots are mean and the green lines show the median. This plot is indicating many cells with zero expression (drop-outs) and comparable average expressions in all the clusters except cluster one as seen in the red box. C) Boxplot showing the expression of MS4A1 in the coverage corrected (CC) data showing similar average expression points (blue dot) and medians (green line) as seen in the red box. Only cluster five is out of the red box and it’s because there are only seven cells in this cluster. D) Boxplot showing the expression of MS4A1 in the data imputed with MAGIC showing varying average expression points (blue dot) and medians (green line). As seen in the plot the means and medians don’t fit in the red box. E) Heatmap showing the expression patterns of the top 5 markers per cluster in the raw data which includes many drop-outs. F) Heatmap showing the expression patterns of the top 5 markers per cluster in the coverage corrected (CC) data showing similar patters to the raw data. G) Heatmap showing the expression patterns of the top 5 markers per cluster in the data imputed with MAGIC showing differing patters to the raw data. As shown, markers for clusters 4, 6 and 8 are not distinguishable, and the markers for cluster 8 are expressed more in cluster 3. This indicates that MAGIC imputation introduces noise to the data.

**Figure 2.**
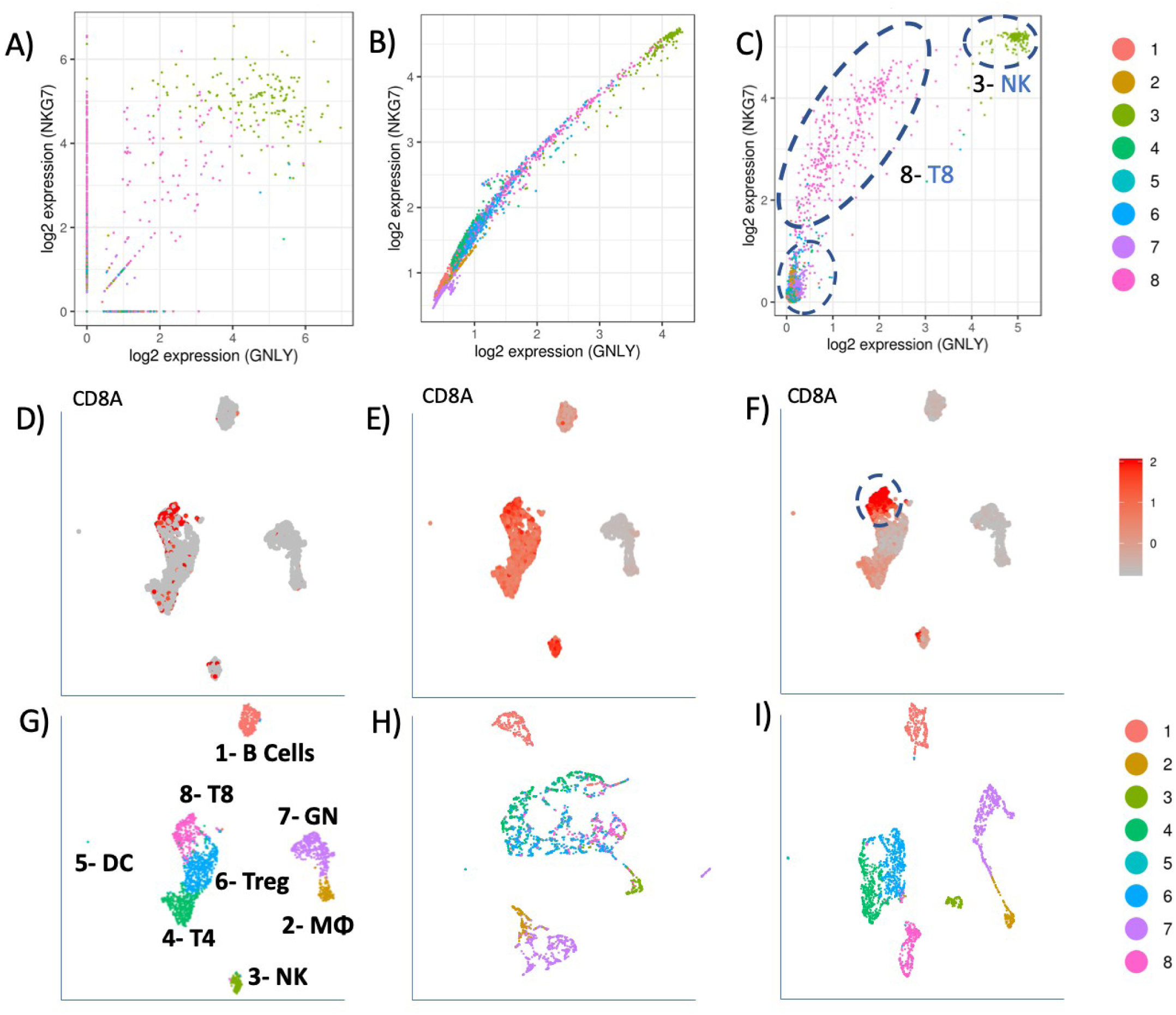
A) Gene-gene correlation for GNLY and NKG7 in the raw data showing many genes with zero counts making it hard to calculate correlation. B) Gene-gene correlation for GNLY and NKG7 in the data imputed with MAGIC showing that NK cells and CD8+ T cells are mixed with other clusters. C) Gene-gene correlation for GNLY and NKG7 in the coverage corrected (CC) data showing that NK cells and CD8+ T cells could be clearly distinguished as they express these genes at much higher levels compared to other clusters. D) UMAP plot showing the expression of CD8A gene in the raw data which includes many drop-outs. E) UMAP plot showing the expression of CD8A gene in the data imputed with MAGIC, showing that it is hard to distinguish CD8-positive cells from other cells. F) UMAP plot showing the expression of CD8A gene in the coverage corrected (CC) data, showing CD8-positve T cells can be clearly distinguished. G) UMAP plot showing the clustering based on non-imputed data. H) UMAP plot showing the clustering based on imputed data using MAGIC. As seen in the plot CD8-positive T cells are mixed with CD4-positive T cells. I) UMAP plot showing the clustering based on coverage corrected (CC) data using iCellR. As seen in the plot CD8-positive T cells are well distinguished from CD4-positive cells also showing more resolution so if clustered again more clusters and details can be seen.

### Combined Coverage Correction normalizes for gene coverage difference across different technologies

We analyzed nine PBMC sample datasets provided by the Broad Institute to detect batch differences [13]. These datasets were generated using varying technologies, including 10x Chromium v2 (3 samples), 10x Chromium v3, CEL-Seq2, Drop-seq, inDrop, Seq-Well and SMART-Seq [13]. Comparing the gene coverages across these technologies shows that combined coverage correction not only corrects for drop-outs but also harmonizes the gene coverages across different technologies (Figure 3).

**Figure 3.**
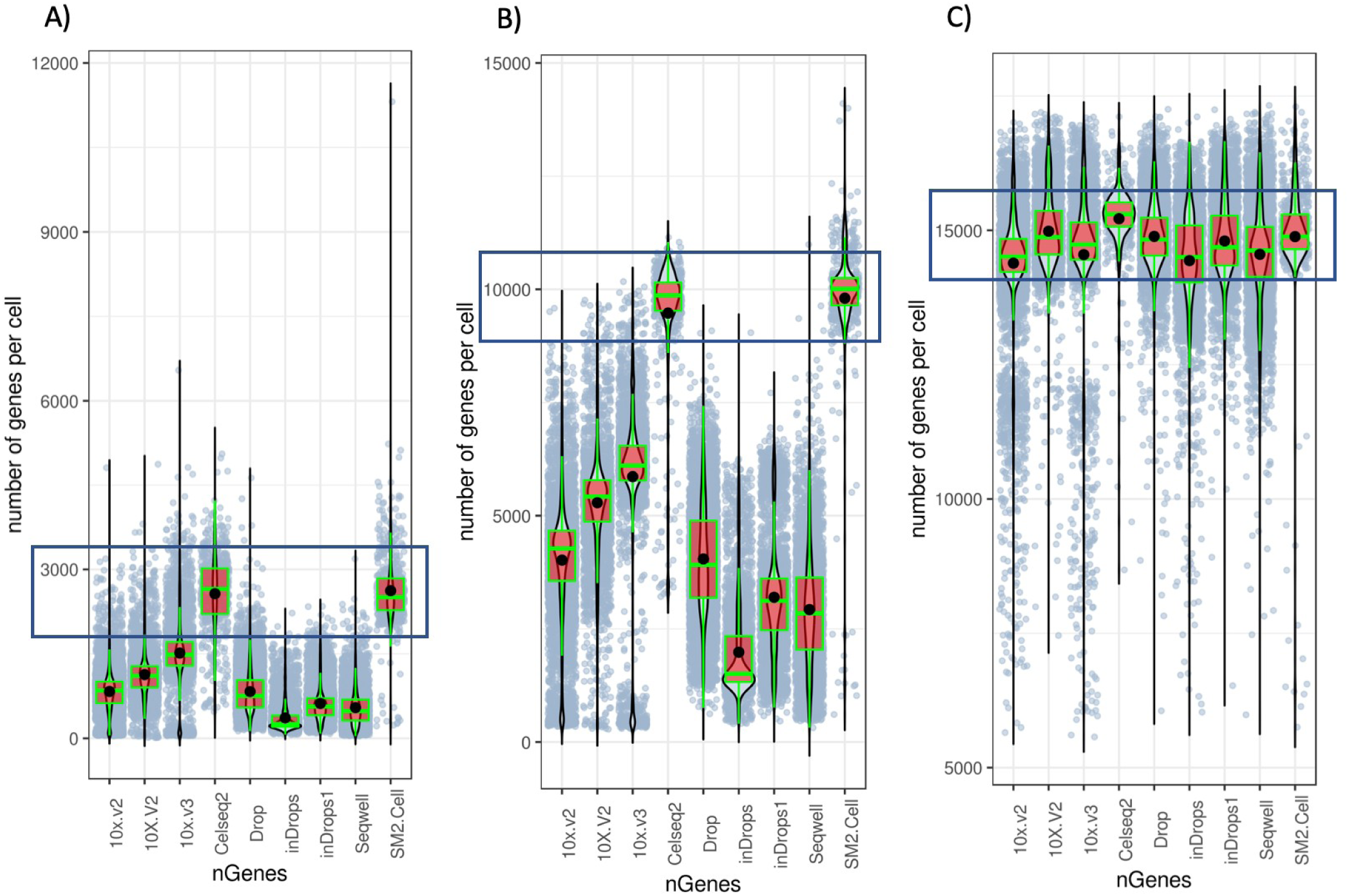
A) Box plots of the gene coverages per cell in the raw data, showing that some technologies have more coverage. B) Box plots of the gene coverages per cell in the coverage corrected (CC) data, showing that the coverages are higher but similar patterns or trends are maintained. For instance, Cel-Seq2 and SMART-Seq2 have the highest coverages. C) Box plots of the gene coverages per cell in the combined coverage corrected (CCCA) data, showing that the coverages across all the technologies are harmonized.

### CCCA and CPCA are both effective at batch alignment

All nine PBMC sample datasets mentioned above were used to perform both CPCA and CCCA alignment. We first performed Principal Component Analysis (PCA) then calculated cell-cell distance for all the cells in the nine samples. Each cell was considered a root cell once and then ten neighboring cells from each sample were found for each root cell. For CCCA, the expression matrix from the 10 samples was used to average the expression values. While for CPCA, the PCs were used. All the averaged values were then applied to the root cell, and this data was then used to perform PCA, T-distributed Stochastic Neighbor Embedding (t-SNE) and Uniform Manifold Approximation and Projection (UMAP). After performing batch alignment using both CCCA and CPCA, we observed that both methods could align the cell types from different technologies correctly (Figure 4 and S3). This was also evident in the heatmaps of marker genes (Figure S1).

**Figure 4.**
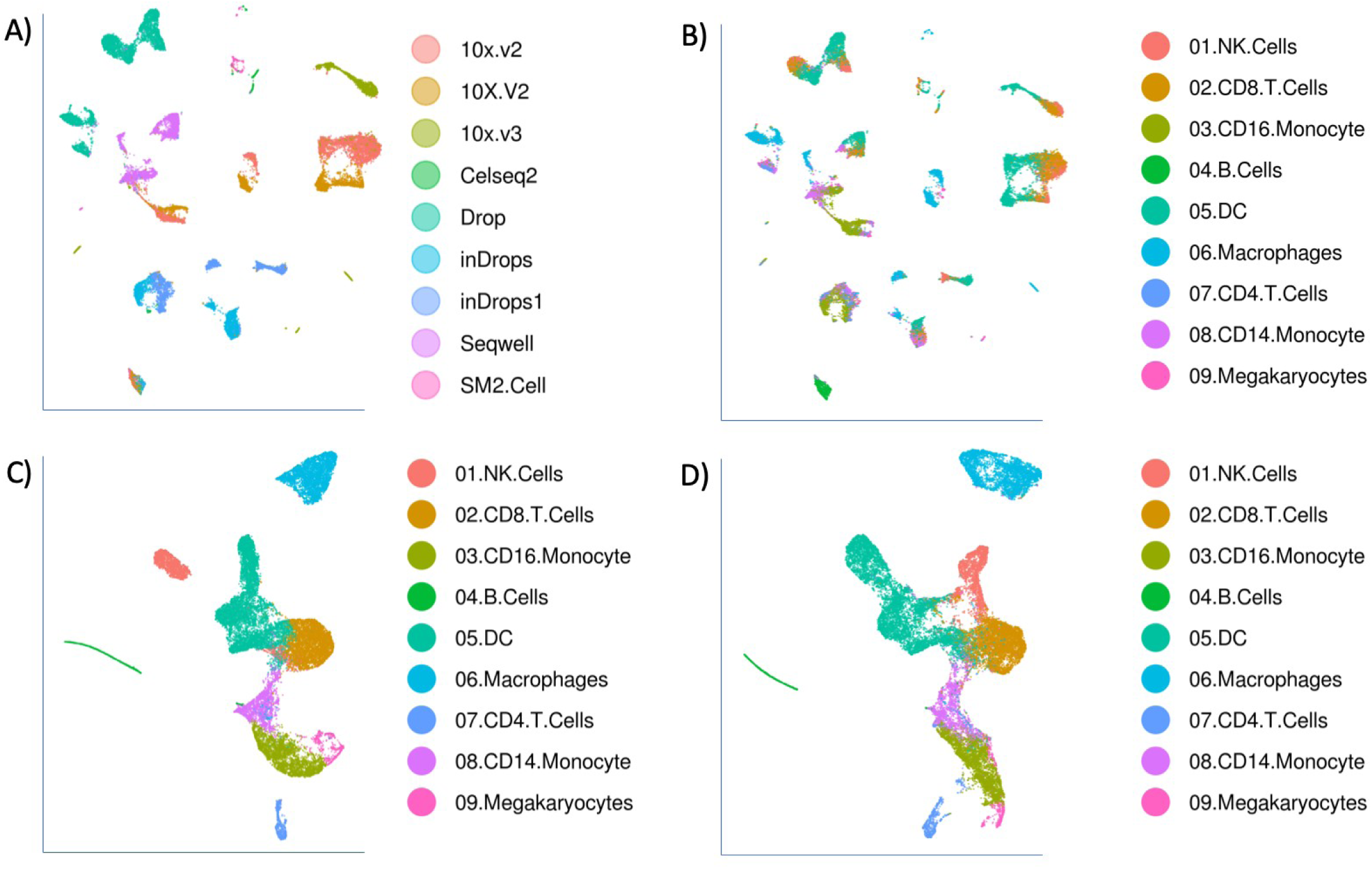
A) UMAP plot of unaligned data, showing the clusters and technologies. B) UMAP plot of unaligned data, showing the cell types from different technologies are not aligned and some colors are seen in multiple clusters. C) UMAP plot of CPCA aligned data, showing that cell types from different technologies are well aligned. D) UMAP plot of CCCA aligned data, showing that cell types from different technologies are well aligned.

## Data Accession and Codes

All the PBMC data we used for batch alignment is available from the Broad Single Cell Portal (https://singlecell.broadinstitute.org/single_cell).

The PBMC data used for coverage correction can be downloaded from the 10x genomics website (https://support.10xgenomics.com/single-cell-gene-expression/datasets).

For connivance all these datasets can also be downloaded from here (https://genome.med.nyu.edu/results/external/iCellR/data/).

All the codes and algorithms are written in R and are available from iCellR package from CRAN (https://cran.r-project.org/package=iCellR).

In addition, the pipelines generating the figures in the results are in supplementary R scripts and the GitHub page for iCellR is (https://github.com/rezakj/iCellR).

## Figures

**Figure S1.**
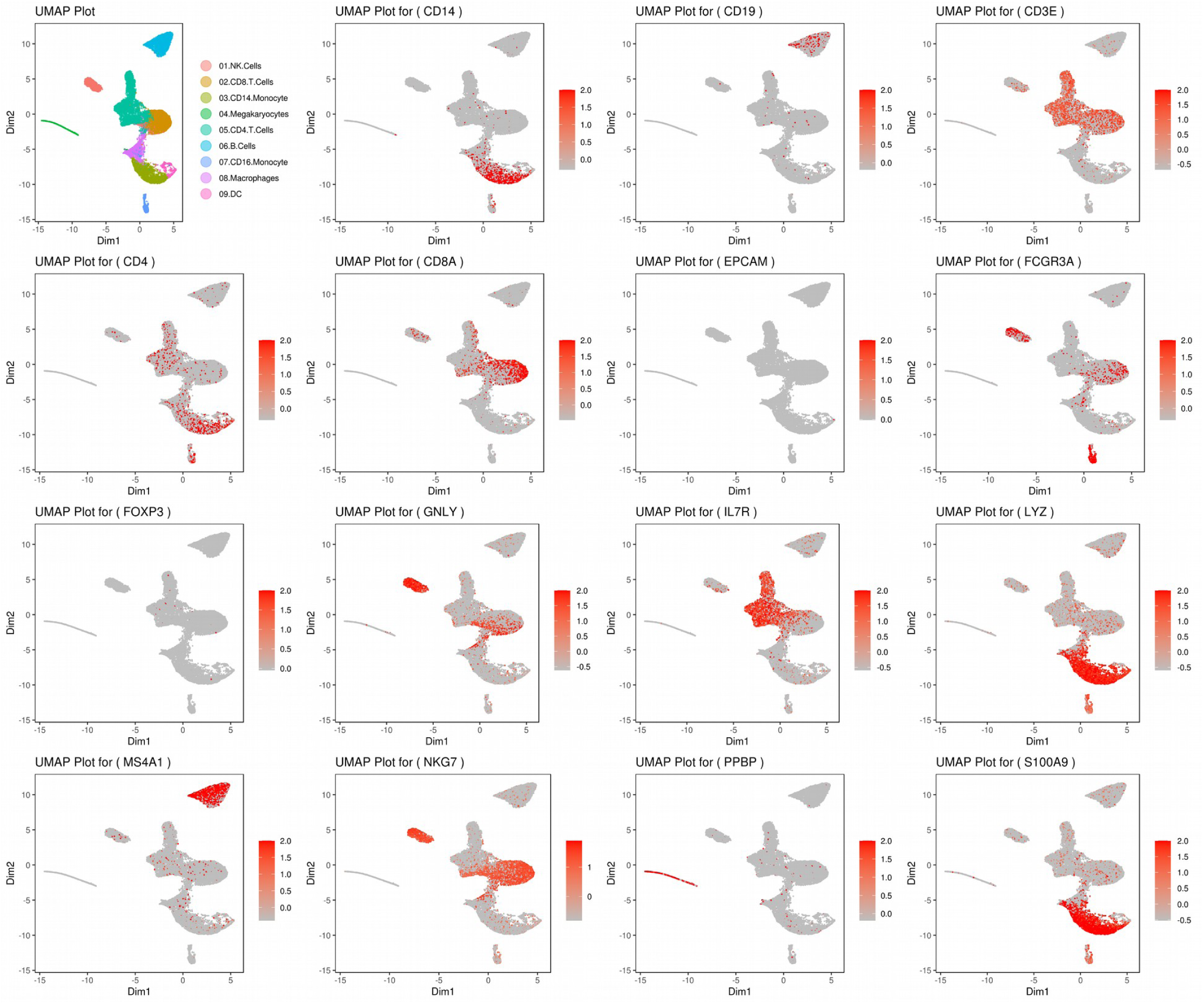
UMAP plots of some gene markers after alignment (CPCA) showing that the samples are properly aligned based on cell types.

**Figure S2.**
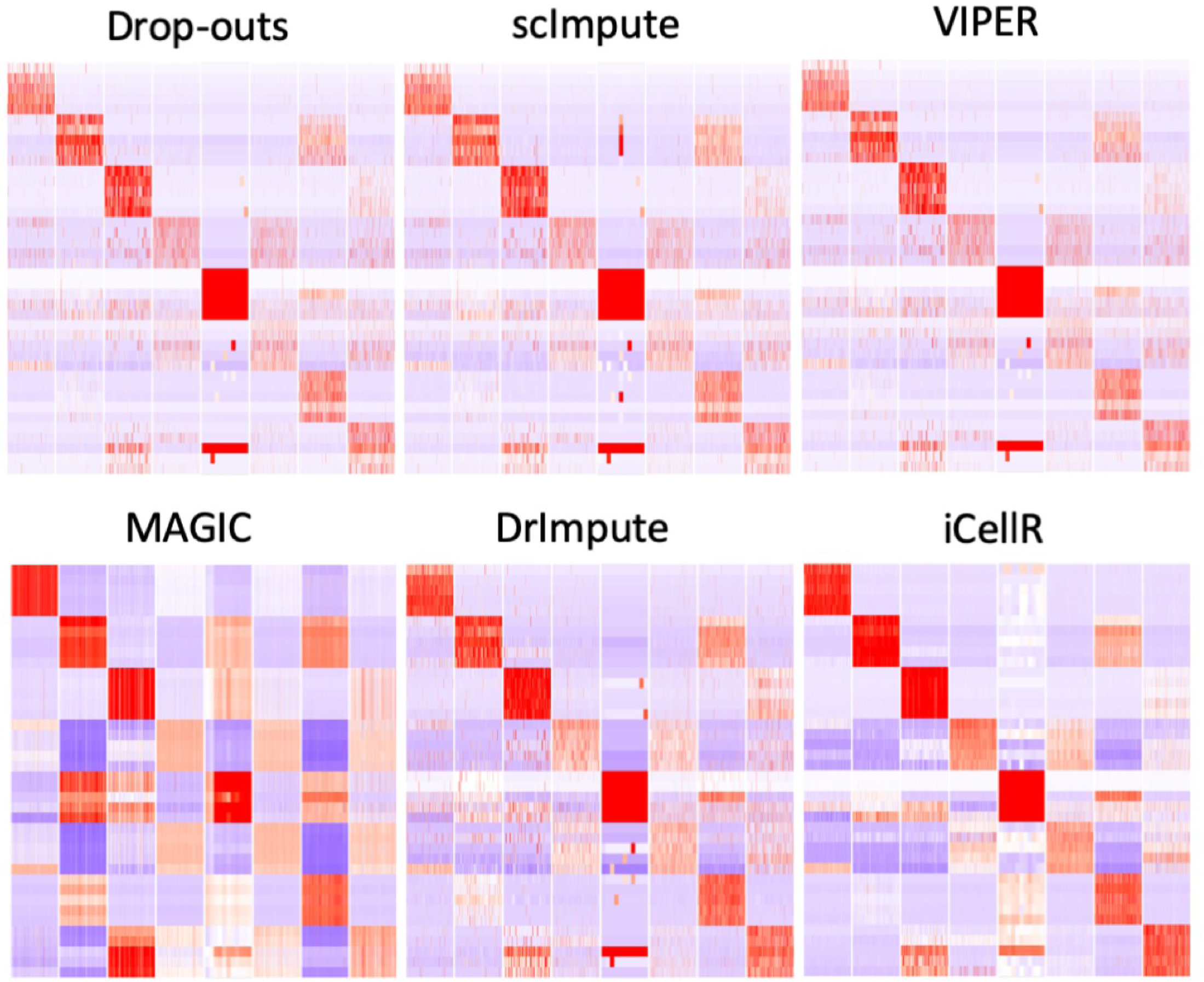
Heatmaps of marker genes before (with high drop-outs) and after imputation (reduced drop-outs). As shown in the figure, iCellR not only fills more drop-outs but also follows similar expression patterns as seen in the original data (Drop-outs).

**Figure S3.**
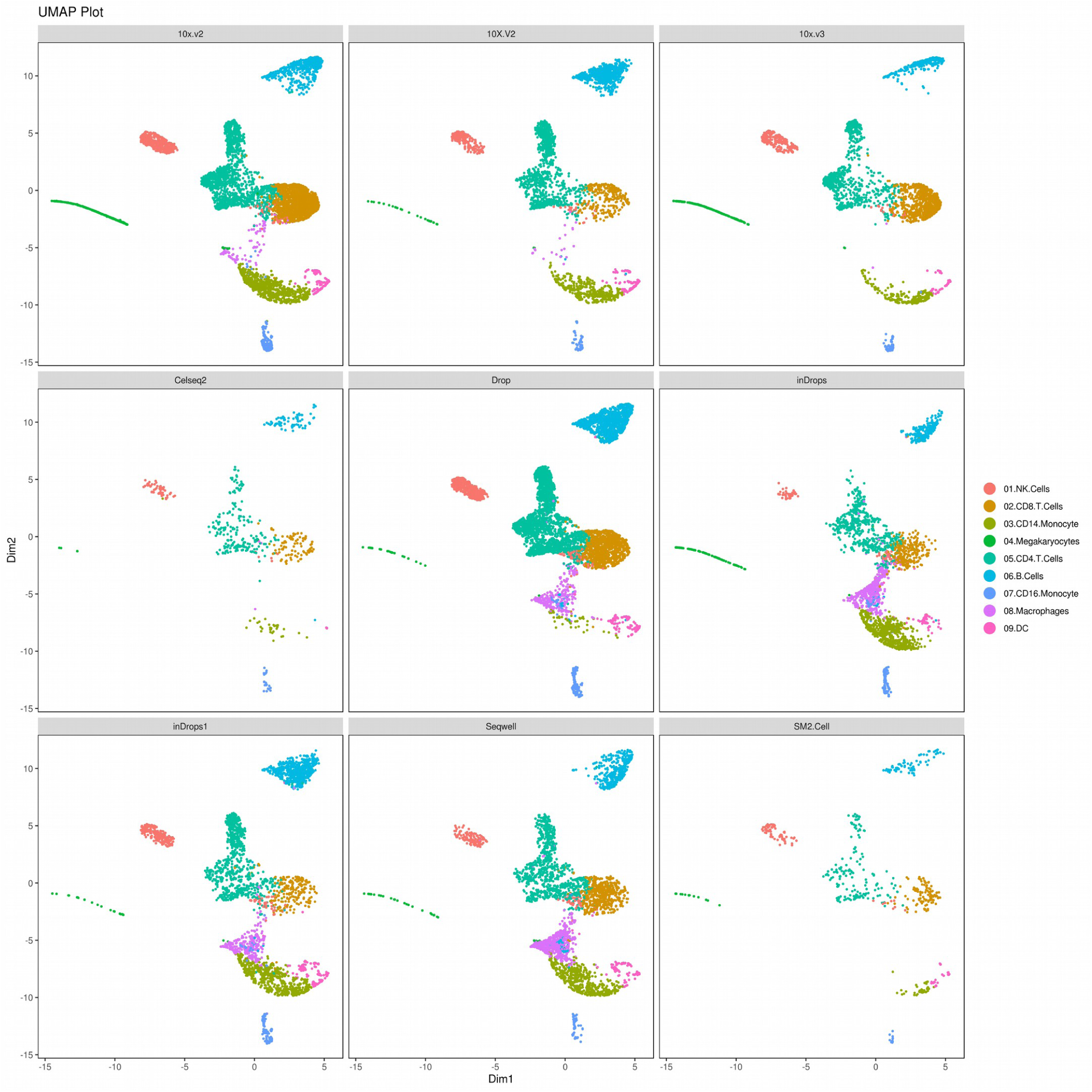
UMAP plots showing the alignments of the nine samples.

## Supplementary Codes 1

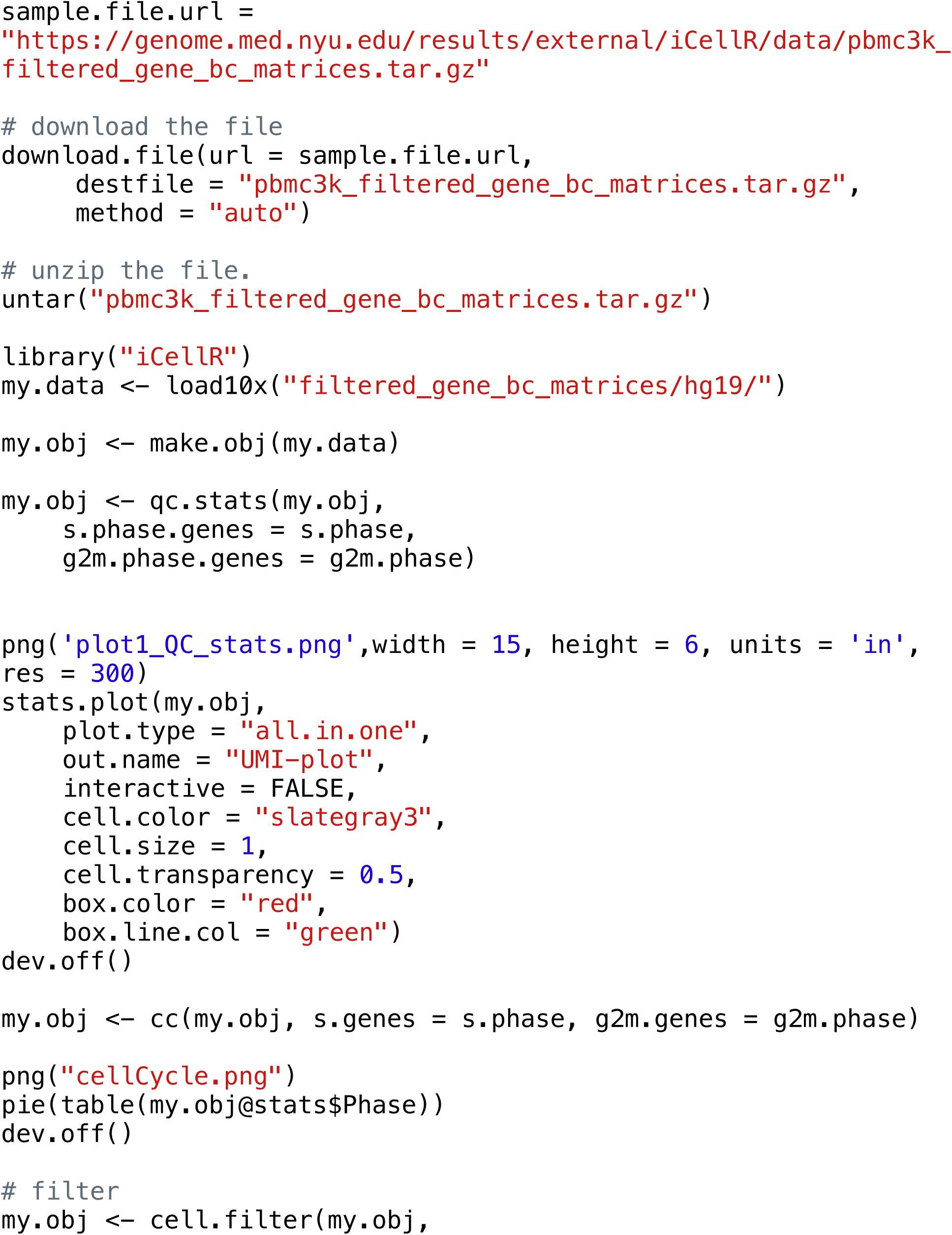
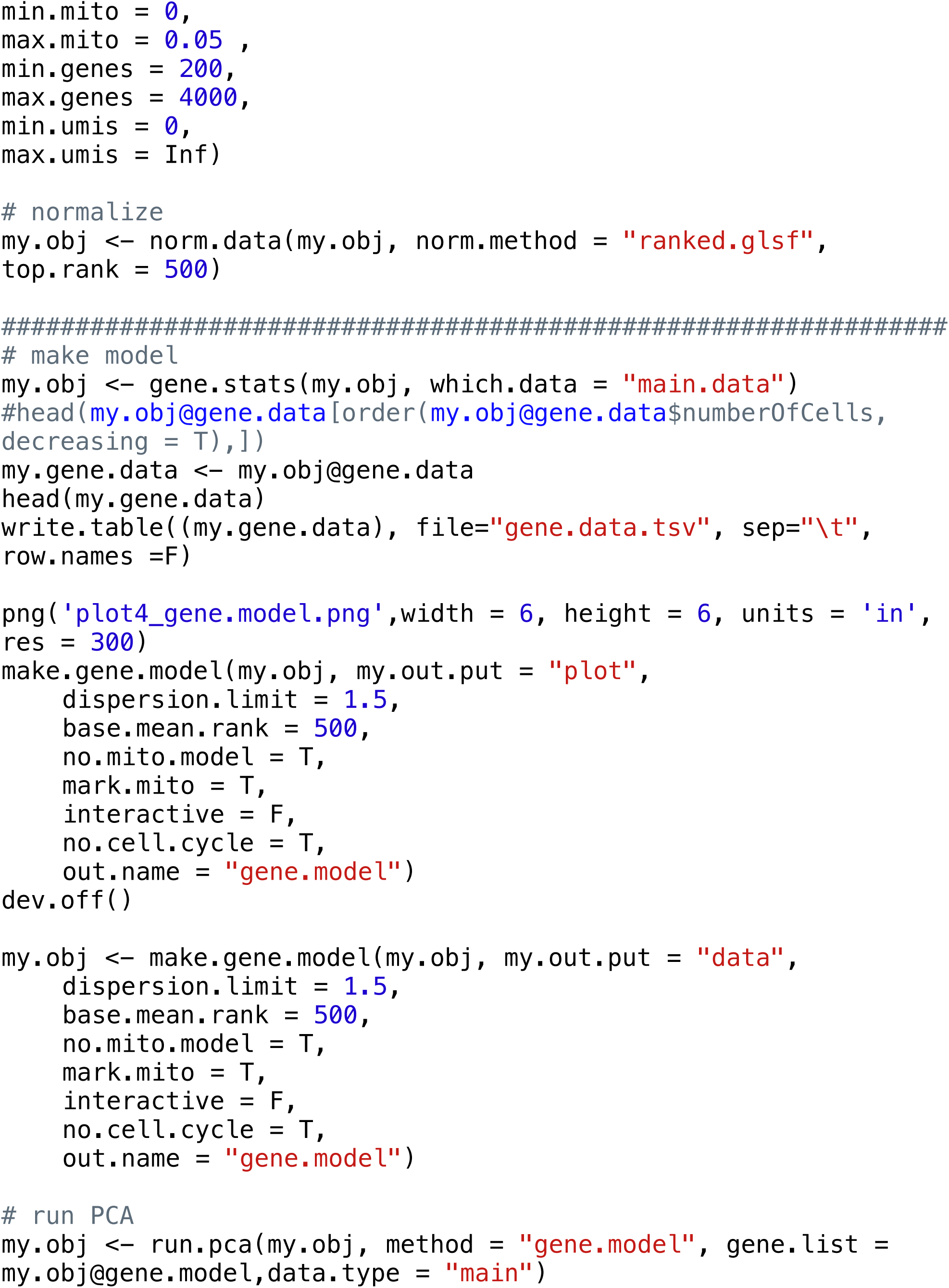
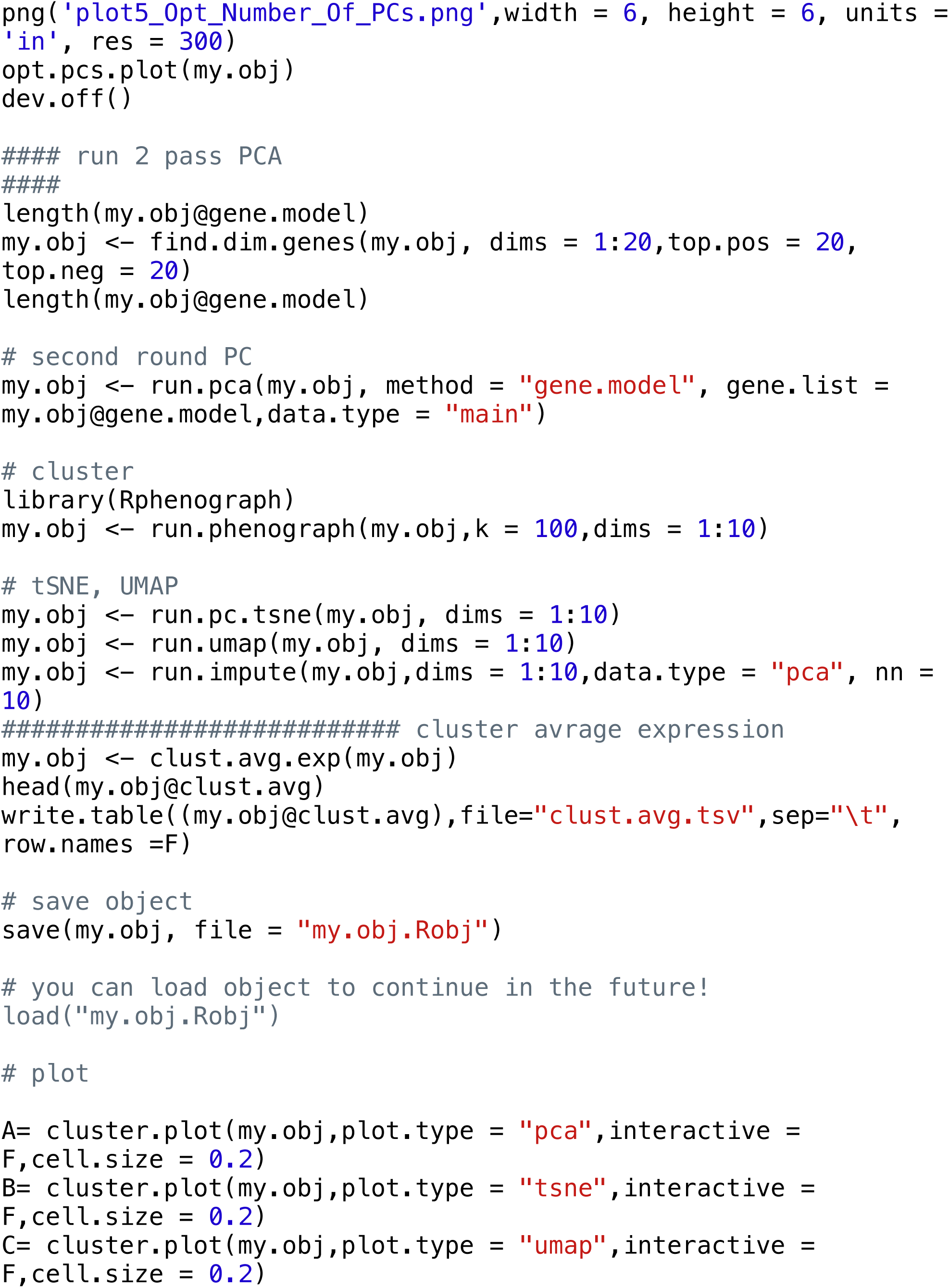
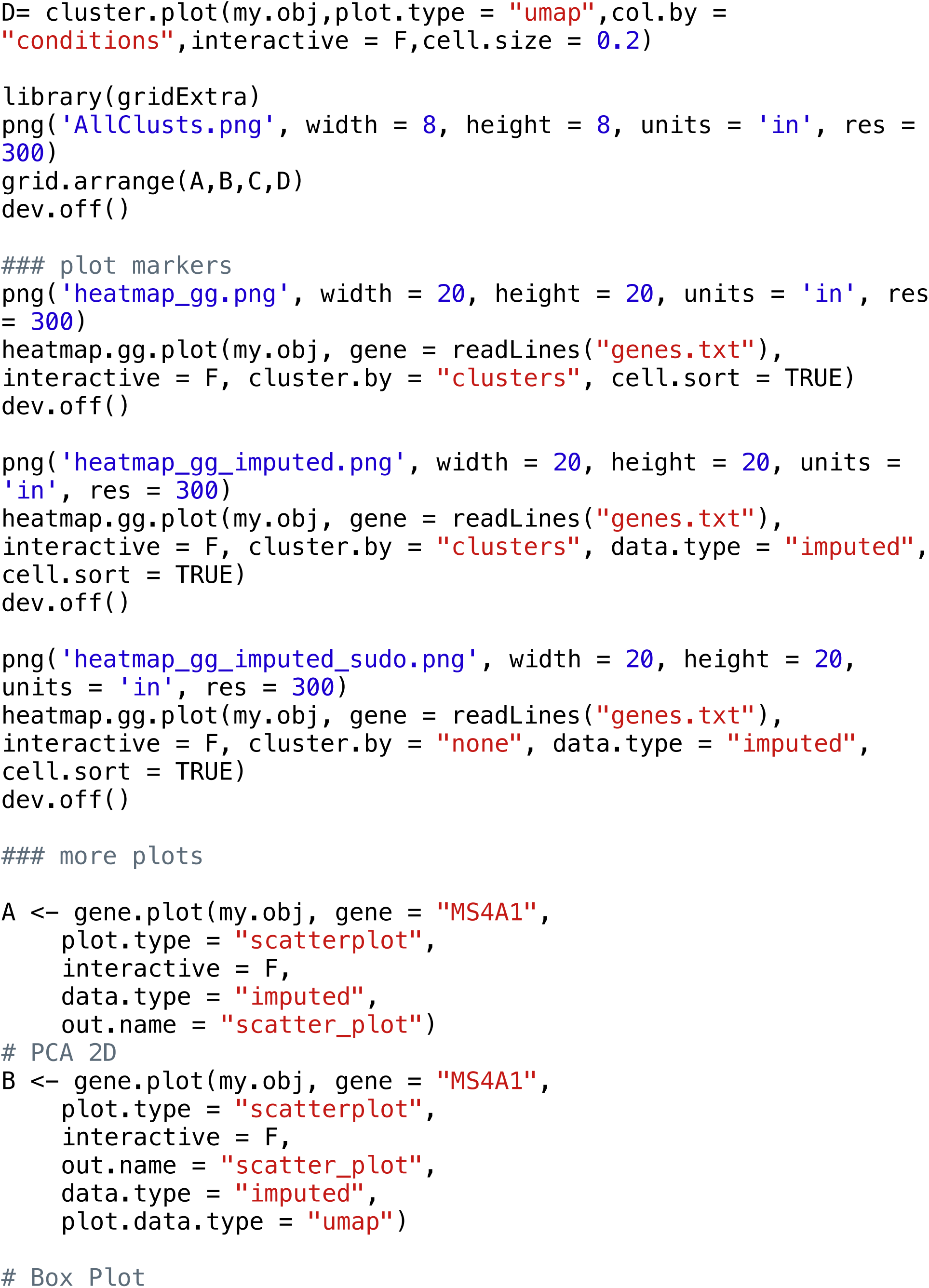
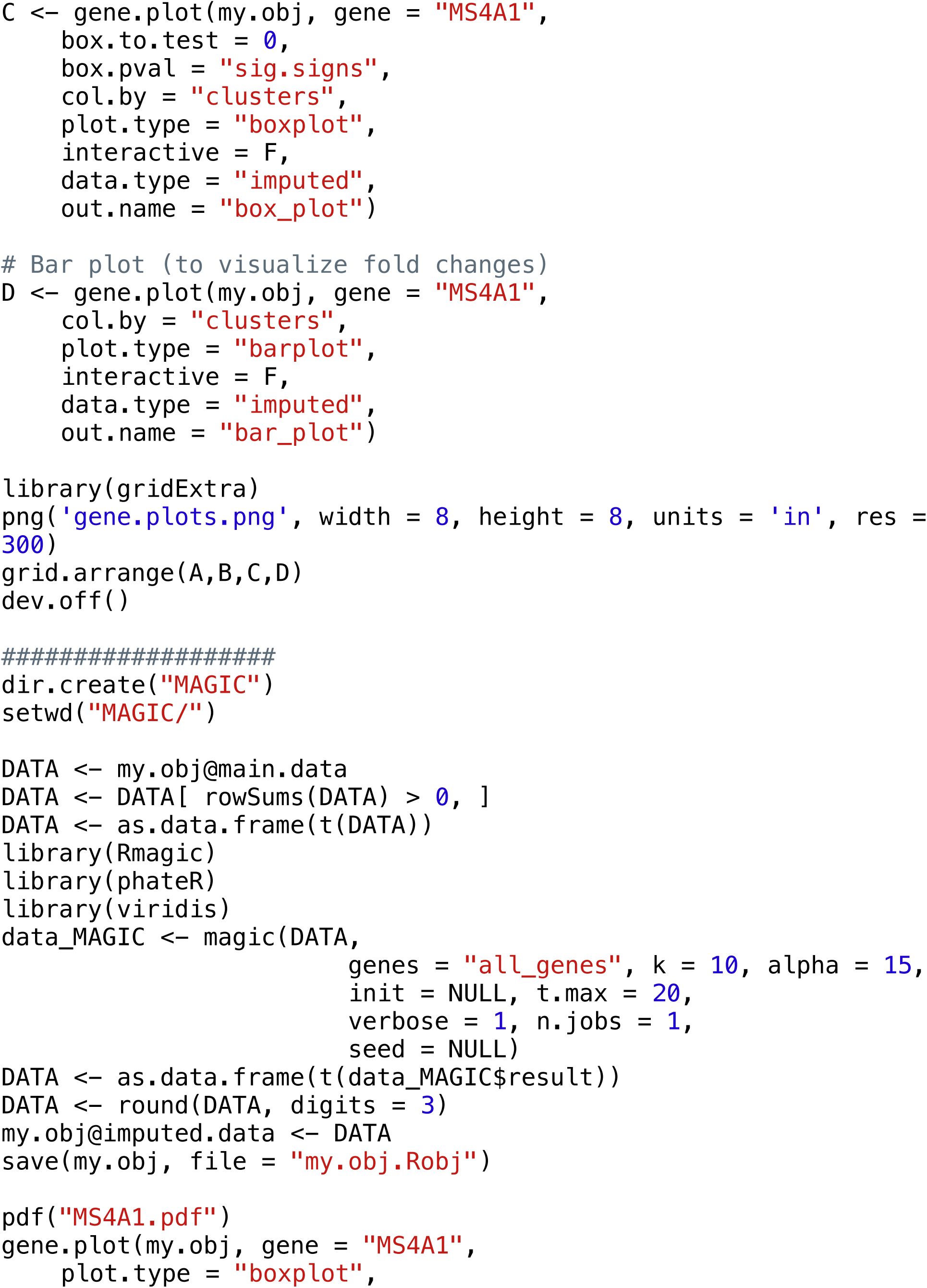
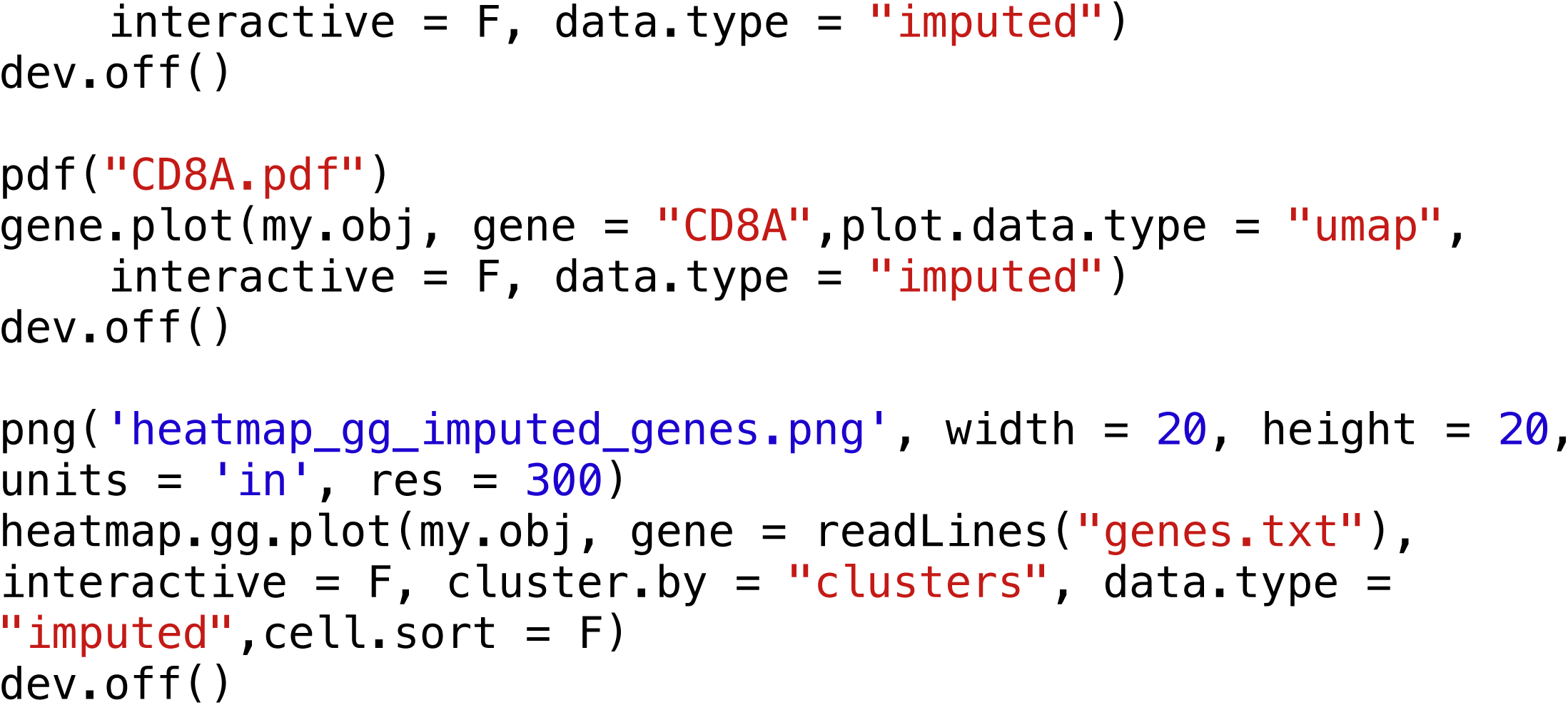

## Supplementary Codes 2

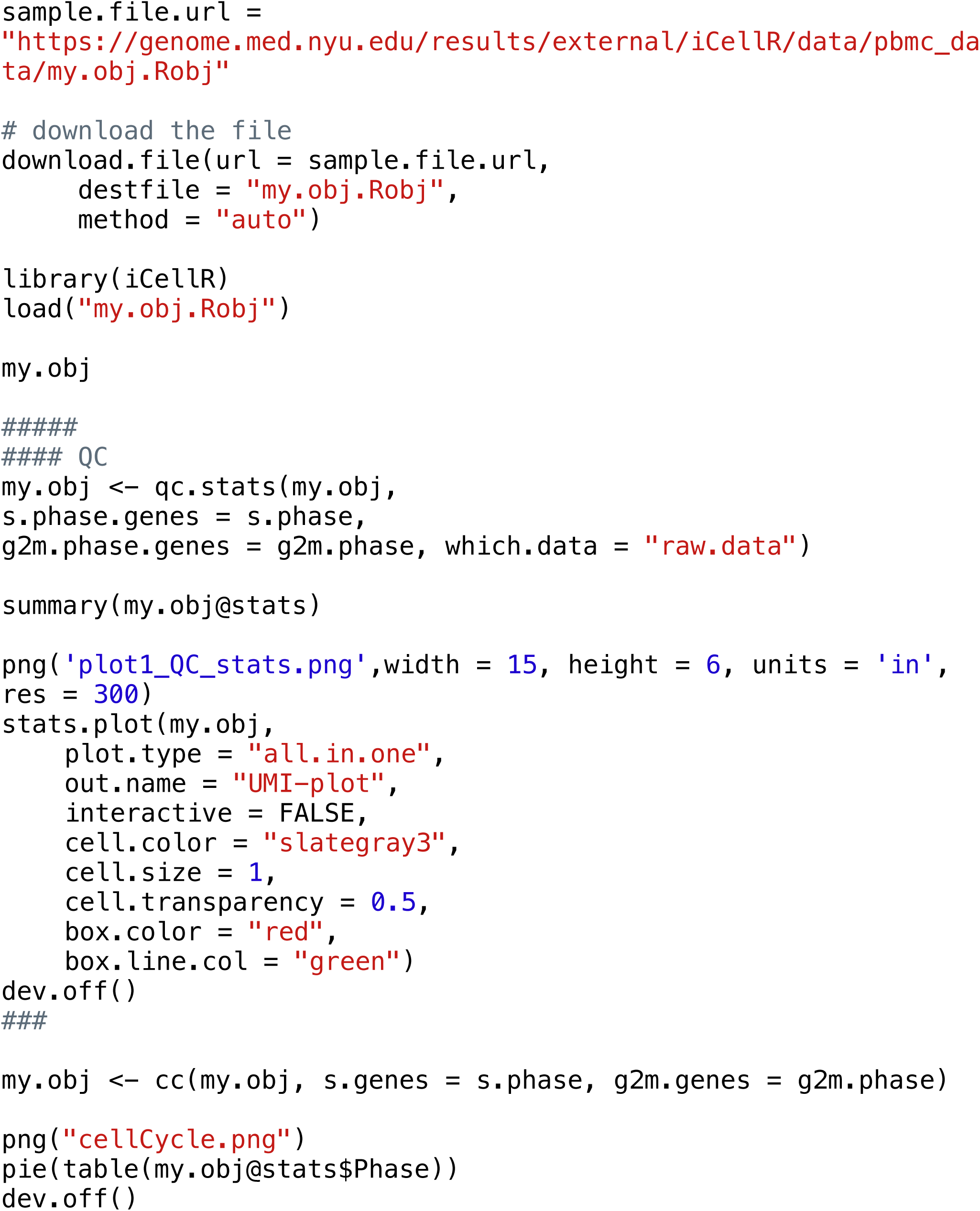
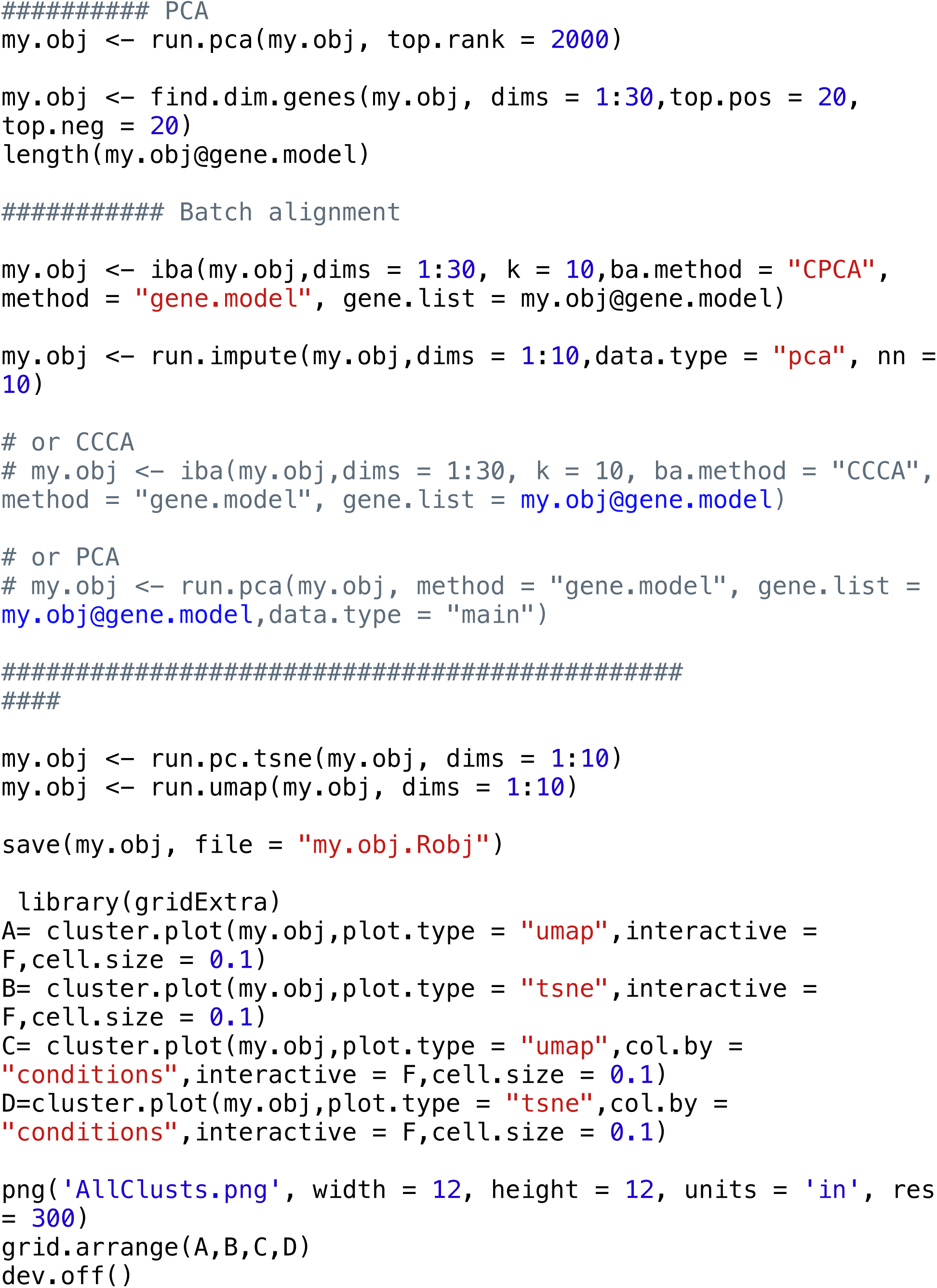
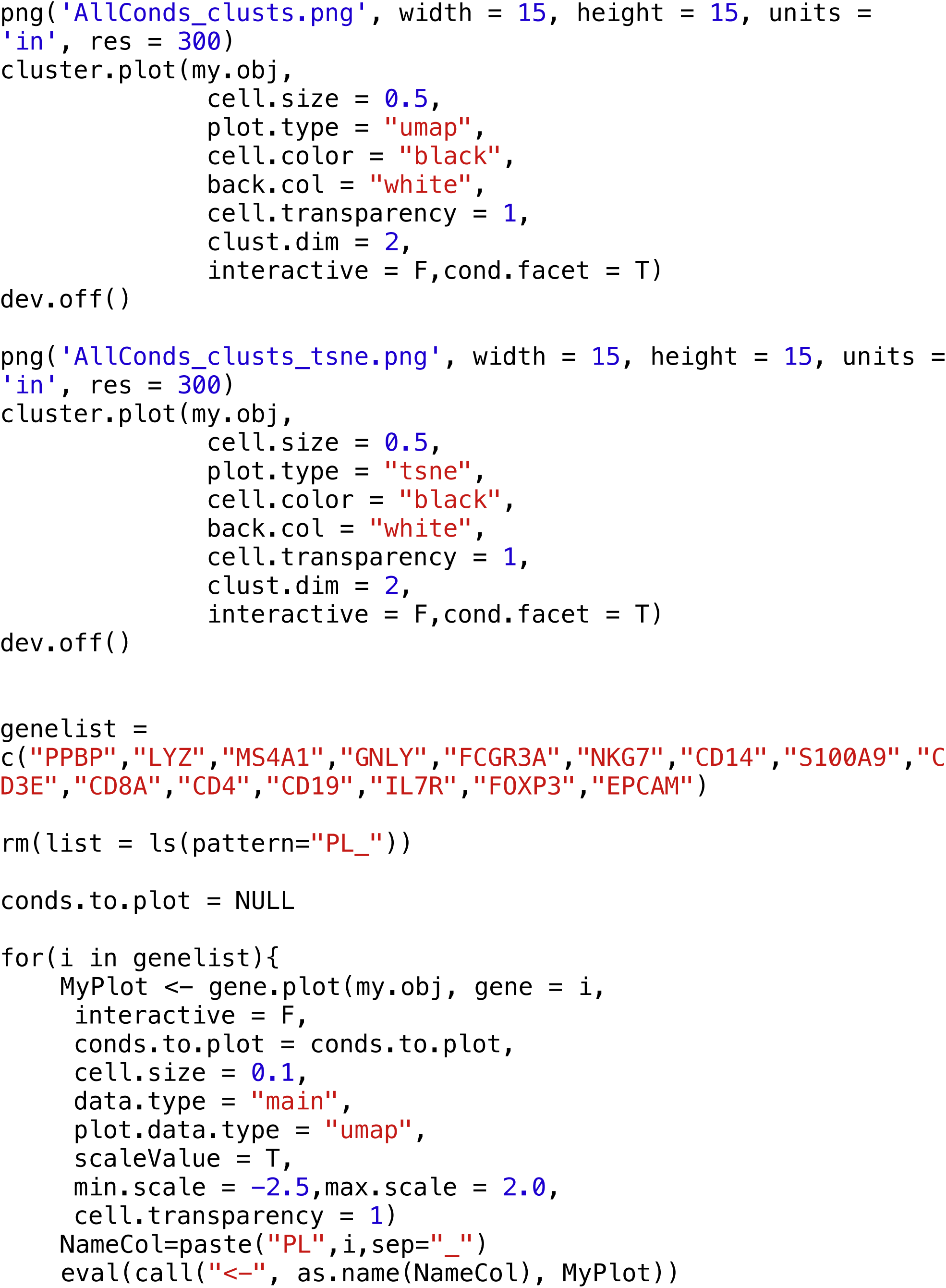
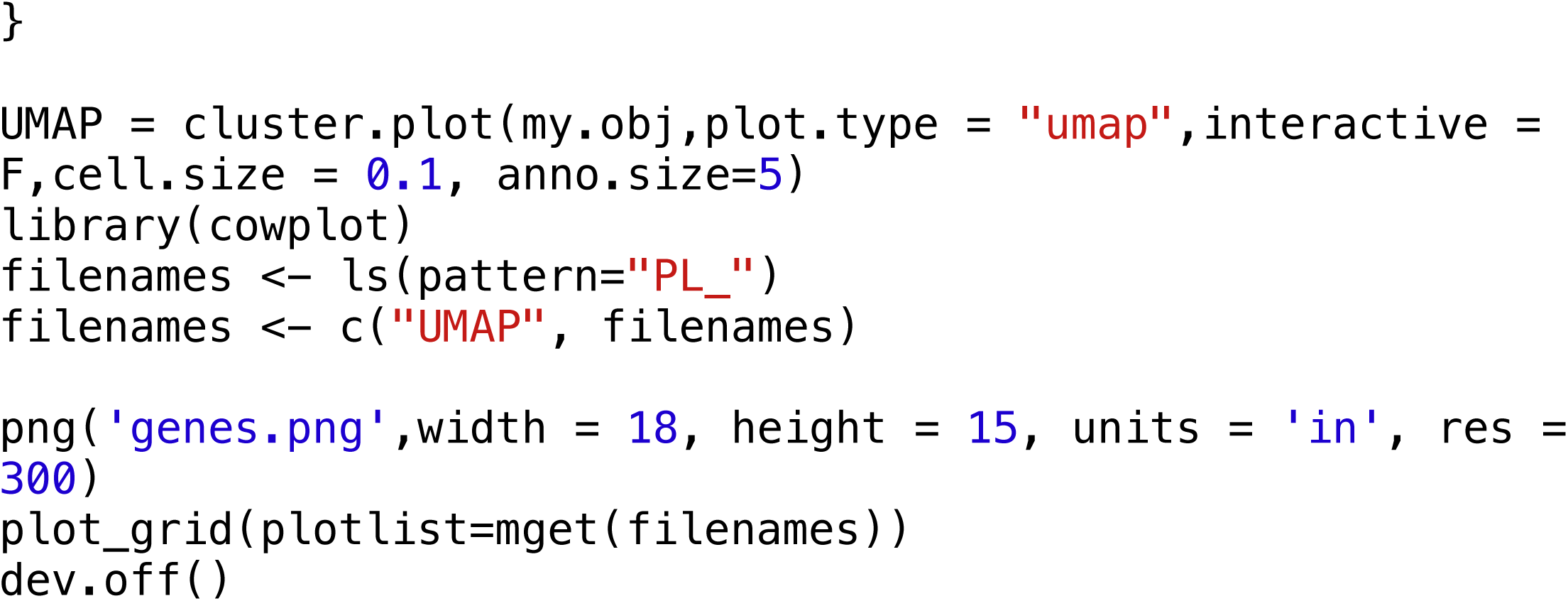

